# Rapid classification of *β*-lactamase activity through AI-based structure prediction and electric field analysis

**DOI:** 10.64898/2026.09.23.753769

**Authors:** Daojiong Wang, Anderson H. Lima, Marc W. van der Kamp

**Affiliations:** School of Biochemistry and Biomedical Sciences, University of Bristol, Bristol BS8 1TD, United Kindgdom; Laboratório de Planejamento e Desenvolvimento de Fármacos, Instituto de Ciências Exatas e Naturais, Universidade Federal do Pará, Rua Augusto Corrêa, 01, 66075-110, Belém, Pará, Brasil

## Abstract

Antibiotic resistance driven by *β*-lactamases has become a major threat to public health. Here, we propose a computational protocol for rapid estimation of *β*-lactamase activity towards carbapenem antibiotics from sequence alone. Starting from AI-predicted acylenzyme complexes, only approximate transition state ensembles are sampled to compute catalytic electric fields, which strongly correlate with experimental activation barriers (*R*^2^ = 0.8). Compared with full quantum mechanics/molecular mechanics (QM/MM) reaction simulations, the protocol substantially reduces the amount of configurational sampling and thus computational cost. Our protocol enables efficient identification of carbapenem breakdown efficiency, further highlighting the important role of electric fields in enzyme catalysis.

---

Antimicrobial resistance has emerged as a major global health threat, causing millions of deaths worldwide^1^. *β*-lactamase enzymes produced by bacteria play a crucial role in the emergence and spread of antimicrobial resistance ^2, 3^. Numerous *β*-lactamase variants with an extended resistance spectrum against *β*-lactam antibiotics further complicate the antimicrobial resistance crisis ^4^. Therefore, efficient estimation of *β*-lactamase activity, ideally based on sequence information only, is urgently needed to rapidly detect resistance-conferring variants and inform antibiotic use and development.

Carbapenem resistance is of particular concern because carbapenems are often regarded as last-resort antibiotics for treating infections caused by multidrug-resistant bacteria ^5^, whereas carbapenem-resistant infections are associated with increased morbidity and mortality ^6^. Many class A *β*-lactamases have carbapenemase activities (including the widely disseminated KPC-2 and its variants) ^7-9^ and they share the same hydrolysis mechanism. Carbapenem (e.g., meropenem) hydrolysis consists of two main stages, acylation and deacylation (**Figure S1**) ^4, 10^. After forming a non-covalent complex with the enzyme, the catalytic residue Ser70 is activated by the general base Glu166 via proton transfer, enabling the hydroxyl oxygen of Ser70 to perform a nucleophilic attack on the electrophilic carbon of meropenem, leading to the formation of an acylenzyme complex. Subsequently, the deacylating water (DW) is activated by Glu166 and performs a nucleophilic attack to hydrolyze the acylenzyme complex. Both stages involve the formation of a tetrahedral intermediate. Previous studies have demonstrated that the energy barrier for the first step of deacylation (acylenzyme complex, AC, to tetrahedral intermediate, TI) correlates with differences in enzyme activity ^11-13^, suggesting that this step is primarily responsible for determining enzymatic activity.

The electric field in the enzyme active site can be a key determinant of catalytic activity ^14, 15^. The tetrahedral intermediate of the deacylation step features a negatively charged carbonyl oxygen (**Figure S1**), which is stabilized by the oxyanion hole of the enzyme ^4^. Changes in the local electric field around the negatively charged carbonyl oxygen can modulate whether a ligand behaves as a substrate or an inhibitor ^16, 17^. Indeed, previous work showed that calculated electric field values can correlate well with experimentally determined carbapenamase activities in class A *β*-lactamases ^18^ (a similar relationship with catalytic activity was recently observed in another enzyme system ^19^), but only when considering transition state or tetrahedral intermediate ensembles. This indicates that such electric field calculations can be used to evaluate *β*-lactamase activity. However, this work relied on computationally expensive extensive sampling of the deacylation reaction free energy landscape, as well as the availability of crystal structures for all enzymes simulated. Alongside the need for detailed structural information, the substantial computational cost of these protocols limits their application for efficient activity assessment of newly emerging *β*-lactamases.

Here, we demonstrate a computational protocol for the rapid estimation of carbapenemase activity of serine-*β*-lactamases, using only protein sequences and acylated antibiotic Chemical Component Dictionaries (CCDs) as input (**Figure 1**). Two groups of class A *β*-lactamases were investigated in this work (as previously ^18^): carbapenemases (KPC-2, NMC-A, SFC-1 and SME-1) ^7, 8, 20-23^, which hydrolyze carbapenems, and non-carbapenemases (BlaC, CTX-M-16, SHV-1, TEM-1, TEM-52, and TEM-116) ^23-26^, which are inhibited by carbapenems. Protenix (Protenix-v1 model) ^27, 28^, the open-source AI-based structure prediction tool, was used to generate the initial acylenzyme complexes. From these complexes, approximate transition state (ATS) structures of the deacylation step were generated via restrained QM/MM minimization at the DFTB2/ff14SB level ^29-31^ (**Figure S2**). To obtain the ATS, restraints were applied to ensure that the deacylating water is close to complete the nucleophilic attack, with the proton transfer to Glu166 approximately halfway. This is based on the transition state location observed in previous mechanistic studies ^12, 13, 18^. The generated ATS structures then underwent QM/MM simulation with the same restraints, using three independent replicas and either 20 ps or 2 ps sampling. This resulted in ATS ensembles of 600 or 60 structures per system for subsequent electric field analysis using FieldTools ^18^ (**Figure 2A**). The calculated electric field was then compared with the experimental activation barriers derived from the experimental deacylation rate constant (*k*_3_) where available, or else from *k*_cat_ (**Table S1**).

**Figure 1.**
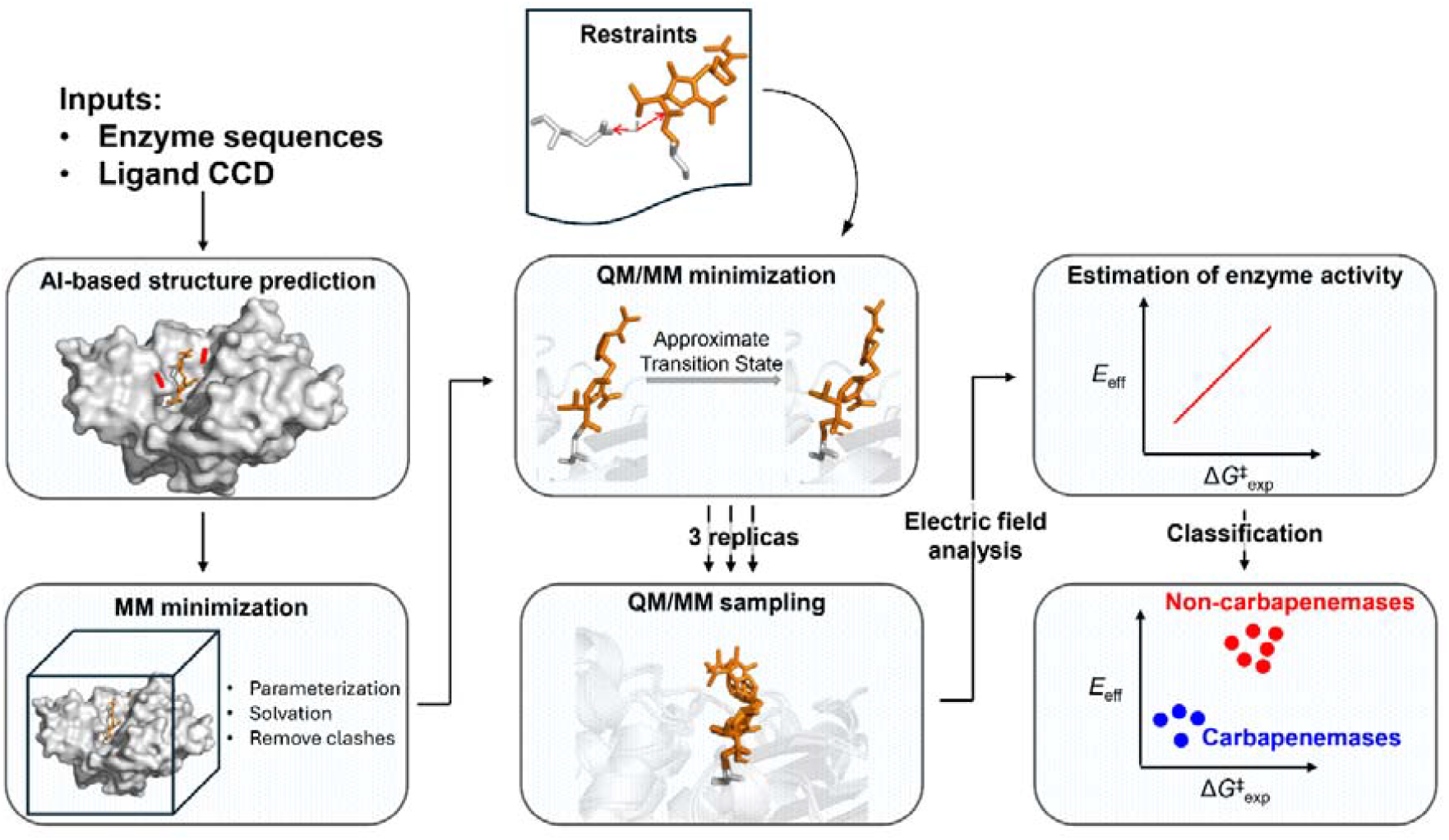
Scheme of the computational protocol for rapidly estimating *β*-lactamase activity.

**Figure 2.**
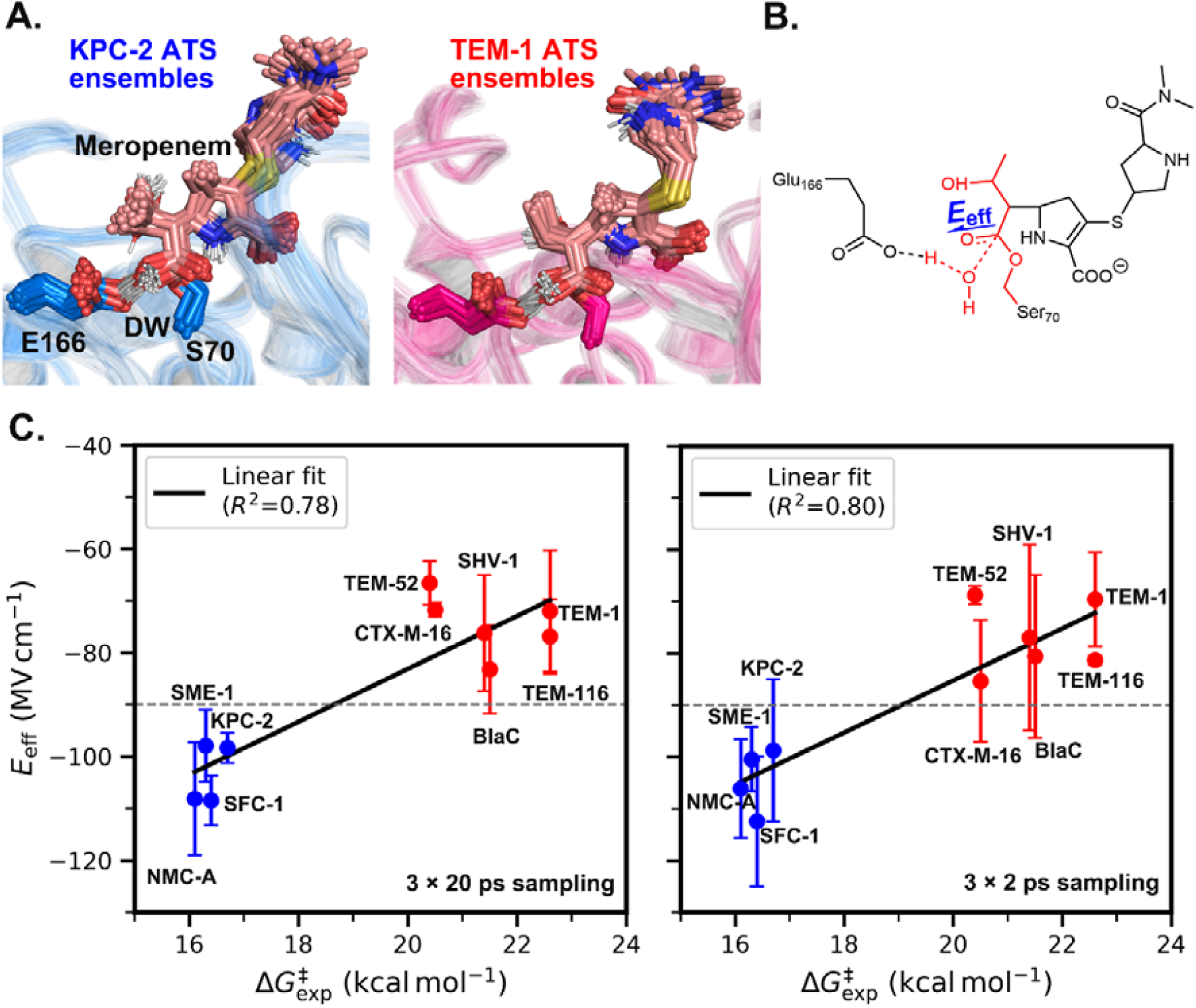
Computed effective electric fields and experimental activation energy barriers. (A) Examples of the sampled approximate transition state (ATS) ensembles (20 frames from each of three replicas, 60 frames per system). Ensembles of carbapenemase KPC-2 (left panel) and non-carbapenemase TEM-1 (right panel) are shown, with the deacylating water (DW), meropenem, and sidechains of the general base Glu166 and catalytic Ser70 shown as sticks. (B) The effective electric field (*E*_eff_) was computed as the projection of the electric field onto the meropenem C=O bond based on ATS sampling of the first step of deacylation. Bonds undergoing changes during the reaction are represented by dashed lines. Atoms excluded from electric field calculations are labelled in red. (C) Calculated total *E*_eff_ (protein and solvent) based on ATS sampling of 20 ps (left panel) and 2 ps (right panel) per replica, with the hydroxyethyl group of meropenem excluded in the calculation. The correlation between *E*_eff_ and experimental energy barrier is shown. Carbapenemases are coloured in blue, whereas the non-carbapenemases are coloured in red. Error bars for *E*_eff_ represent the standard deviation of the mean across three replicas.

The hydroxyethyl group of carbapenems has been observed to adopt three different conformations in crystal structures of class A *β*-lactamases ^13^. Based on the dihedral angle defining the orientation of the hydroxyethyl group (**Figure S3**, first panel), the conformations can be classified as I (∼50°), II (∼200°), and III (∼290°). Previous simulations showed that these three different conformations could be sampled and led to differences in enzyme activity in both class A and class D *β*-lactamases ^13, 32^. Here, our Protenix predicted acylenzyme complexes of class A *β*-lactamases with meropenem also show distinct conformations of the hydroxyethyl group (**Figure S3**). Interestingly, the hydroxyethyl group adopts conformation II in all four carbapenemase acylenzyme complexes (KPC-2, NMC-A, SFC-1 and SME-1), whereas in non-carbapenemase complexes it adopts conformation III in BlaC, a ∼360° orientation in CTX-M-16, TEM-1, TEM-52, and TEM-116, and a ∼90° orientation in SHV-1.

The conformations of the hydroxyethyl group in the sampled ATS ensembles show a similar trend: conformation II is dominant in all four carbapenemases, whereas conformation III is dominant in all non-carbapenemases except TEM-52, in which conformation I is the major conformation (**Figure S4**). The similarity is likely due to limited sampling (20 ps simulation per replica), with the sampled hydroxyethyl group conformations being highly dependent on the initial predicted acylenzyme complexes.

The effective electric field (*E*_eff_) was defined as the component of the electric field aligned along the meropenem C=O bond (**Figure 2B**). To minimize contributions arising from the reaction and substrate conformational changes, the same “reactive” part of the substrate defined in the previous work ^18^, including the deacylating water, the Ser70 sidechain (including Cα), the carbonyl group and adjacent carbon atoms (with their attached hydrogens), was excluded from the electric field analysis. Initially, the hydroxyethyl group was included (as previously). Based on this, the calculated total *E*_eff_ (protein and solvent) based on ATS ensembles shows a weak correlation with experimental energy barriers 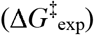 (**Figure S5**, *R*^2^ = 0.44). However, the observed trend is still consistent with the expectation that a more negative *E*_eff_ along the C=O bond would stabilize the transition state and thus lower the energy barrier ^18^. This weak correlation (compared to *R*^2^ = 0.68 in ref. ^18^) may result from differences in the sampled hydroxyethyl group conformations. Notably, TEM-52 is the main outlier in the overall regression, with by far the least negative *E*_eff_ (**Figure S5**). It is also the only enzyme where the hydroxyethyl group is sampled in conformation I (**Figure S4**, almost exclusively). This conformation was indicated to lead to low activation barriers in both class A and D *β*-lactamases ^13, 32^, but this is expected to be due to facilitating the proton transfer from the deacylating water to the active site base. This effect will thus not be captured by the *E*_eff_ on the meropenem C=O bond. Furthermore, possible hydroxyethyl conformations are not sufficiently explored in the short sampling performed here.

To avoid the potentially large influence of the hydroxyethyl group conformation on our electric field calculations (due to its close proximity to the meropenem C=O bond), we excluded this group (**Figure 2B**). The resulting *E*_eff_ values show a significantly stronger correlation with Δ*G*^‡^_exp_ when using the same trajectories (**Figure 2C**, left panel, *R*^2^ = 0.78). When only the first 2 ps of each trajectory are used to reduce the computational cost further, the correlation is similar (*R*^2^ = 0.80), although the associated standard deviation is larger (**Figure 2C**, right panel). The average *E*_eff_ value for each system is also similar (largest change for CTX-M-16), suggesting that even very short sampling is sufficient for carbapenemase activity estimation. Compared with a previous study that used extensive 2D umbrella sampling to scan the full free energy surface (using 10 replicas per system with over 3.8 ns of QM/MM simulation per replica) ^18^, the present computational protocol significantly reduces the cost, only requiring 25 or 7 ps of simulation for each of 3 replicas (5 ps relaxation and 20 or 2 ps sampling), while providing an excellent correlation. The *E*_eff_ values also differ significantly between carbapenemases and non-carbapenemases (independent two-sample, two-tailed *t* test, p < 0.001). The backbone amide of oxyanion-hole residue 237 (TEM-1 numbering) forms a hydrogen bond with the carbonyl oxygen, which stablizes the transition state and tetrahedral intermediate of deacylation. Previous work demonstrated that the highest correlation was obtained from the electric field contribution of the oxyanion hole residue 237 (*R*^2^ = 0.86) ^18^. We confirm that this is also the case here (**Figure S6**, *R*^2^ = 0.89 with either 20 ps or 2 ps sampling). These results further indicate the importance of the oxanion hole in enzyme catalysis. Not only the enzyme activity of serine *β*-lactmases, but also other serine hydrolases and proteases could potentially be estimated rapidly through analyzing the electric field contributions of the residue(s) forming the oxyanion hole on (approximate) transition state or tetrahedral intermediate ensembles.

Notably, the ATS ensembles were generated using the same reaction coordinates for all enzymes, rather than enzyme-specific coordinates as adopted in previous study ^18^. Although this leads to a less rigourous representation of the transition state compared to those obtained previously (based on the free energy surface of the deacylation reaction), chemically similar ensembles are compared in each enzyme, and a strong correlation of *E*_eff_ with experimental data is obtained. Here, partial charges assigned to the QM region atoms from the MM force field were used in the calculation of the electric field, implying that other semi-empirical methods or indeed machine-learning potentials ^33^ (even if they do not result in accurate transition state energies) can also be employed. The deacylation ATS-derived environmental electric fields separate carbapenemases from non-carbapenemases and correlate well with free energy barriers derived from experimental meropenem turnover, further supporting that deacylation is likely rate-limiting for carbapenem hydrolyisis in class A *β*-lactamases, consistent with previous experimental and QM/MM umbrella sampling studies ^11, 12, 34^. Similarly, a weaker correlation is observed between protein effective electric fields (excluding solvent) and experimental activation energy barriers (**Figure S7**, *R*^2^ = 0.52), in agreement with previous work ^18^, indicating that the active site solvent contributes to differences in the carbapenemase activity ^32, 35, 36^. However, exact solvent positions (e.g. as determined by X-ray crystallography) are not required, as our protocol applies standard (crude) water placement.

The same protocol was also used to sample acylenzyme complex ensembles and analyze the electric field. The conformational sampling of the hydroxyethyl group in acylenzyme complexes is comparable to that in ATS ensembles. Conformation II remains dominant in carbapenemases except KPC-2, whereas conformation III is the major conformation in all remaining enzymes (**Figure S8**). No significant sampling of conformation I was observed. The correlation between acylenzyme total *E*_eff_ (with hydroxyethyl group excluded) and experimental data (**Figure S9**, *R*^2^ = 0.56 for protein and solvent, *R*^2^ = 0.37 for protein only) is significantly weaker than that obtained with ATS ensembles. This demonstrates that the (approximate) transition state provides a more appropriate configuration for predicting differences in energy barriers, consistent with the crucial role of transition state stabilization in enzyme catalysis. Consistent with this, the total and protein *E*_eff_ values from carbapenemase ATS ensembles are generally more negative than those obtained from the corresponding acylenzyme complexes (**Table S1**), illustrating that the transistation state stabilization is reflected in the *E*_eff_.

Due to limited sampling, the accuracy of our protocol will depend on the quality of the initial AI-predicted structures. Only crystal structures of three simulated meropenem acylenzyme complexes (BlaC, SHV-1, SFC-1 E166A, and SFC-1 S70A; **Table S2**) were included in model training of the method used (Protenix-v1 PDB training set cutoff was 30 September 2021). Several relevant other carbapenem (imipenem) acylenzyme complexes (TEM-1 N132A, TEM-116, KPC-2 F72Y, SME-1; **Table S2**) were also available. No carbapenem acylenzyme complexes with CTX-M-16, TEM-52, or NMC-A were included in the training set, indicating that AI-based structure prediction based on other data available is sufficient to generalize to “new” enzyme-carbapenem complexes for activity estimation by our protocol.

In summary, our work presents an efficient computational protocol for evaluating *β*-lactam hydrolysis by *β*-lactamases. By using AI-based structure prediction to generate acylenzyme complexes for ten different class A *β*-lactamases and sampling only approximate transition state structures of the first step of deacylation, the computed total effective electric field (*E*_eff_) along the meropenem C=O bond shows strong correlation with experimental data, with two distinct populations corresponding to carbapenemases and non-carbapenemases. For the first time, carbapenemase activity can thus be rapidly estimated from sequence information alone. Our results demonstrate that electric field calculations capture differences in enzyme activity arising from subtle mutations and conformational variations. With the rapid rise of antibiotic resistance, this protocol may help identify *β*-lactamase activity spectra, thereby facilitating the selection of appropriate therapeutic strategies as well as future rational design of antibiotics to combat antibiotic resistance.

## Data Availability

The complete protocol, together with worked example, will be made freely available online upon publication.

## Author Contributions

M.W.v.d.K. and A.H.L. conceived the work. D.W. performed all simulations. D.W., A.H.L. and M.W.v.d.K. performed analysis and interpretation of results. M.W.v.d.K. supervised D.W. D.W. and M.W.v.d.K. wrote the manuscript draft, with revisions and approval by all authors.

## Notes

The authors declare no competing financial interest.

## Acknowledgements

D.W. and M.W.v.d.K. thank the GW4 BIOMED2 DTP for DW’s studentship, grant MR/W006308/1 awarded to the Universities of Bath, Bristol, Cardiff, and Exeter from the Medical Research Council (MRC)/UKRI. A.H.L. and M.W.v.d.K. thank the Royal Society for support (Newton International Fellowship to A.H.L, NIF\R1\221443). A.H.L. acknowledges support from the Conselho Nacional de Desenvolvimento Científico e Tecnológico (CNPq; Grant No. 303869/2026-7). All simulations and electric field analysis were conducted using the facilities of the Advanced Computing Research Centre at the University of Bristol (https://www.bris.ac.uk/acrc/).

