## Supporting Information for "Rapid classification of *β*-lactamase activity through AI-based structure prediction and electric field analysis"

This Supporting Information contains:

– Detailed description of the methods applied in this work (Methods);
– Reaction mechanism of meropenem hydrolysis by class A $\beta$-lactamases (Figure S1);
– Description of the QM region used in the QM/MM samplings (Figure S2);
– Visualization of different meropenem 6$\alpha$-hydroxyethyl orientations in Protenix predicted acylenzyme complexes (Figure S3);
– Meropenem 6$\alpha$-hydroxyethyl orientation sampling of approximate transition state and acylenzyme QM/MM samplings (Figure S4, Figure S8);
– Total and protein effective electric fields from approximate transition state and acyleznyme samplings with the 6$\alpha$-hydroxyethyl group included or excluded (Figure S5, Figure S7, Figure S9, Table S1);
– Oxyanion-hole residue 237 denoting effective electric fields from approximate transition state sampling (Figure S6);
– Availability of carbapenem acylenzyme complexes with studied class A $\beta$-lactamases in the PDB database before the Protenix-v1 training set cutoff (Table S2).

### Methods

#### *Acylenzyme complex prediction*

Ten class A $\beta$-lactamases investigated in a previous study [1] were selected in this work. These enzymes can be classified into two groups: carbapenemases (KPC-2, NMC-A, SFC-1 and SME-1) [2-7], which efficiently hydrolyse most carbapenems, and non-carbapenemases (BlaC, CTX-M-16, SHV-1, TEM-1, TEM-52, and TEM-116) [6, 8-10], which are inhibited by carbapenems. The enzyme sequences were retrieved from the Entrez database at NCBI (https://www.ncbi.nlm.nih.gov/protein/). GenPept IDs for enzymes are: AAB07556 for BlaC; AAK32961 for CTX-M-16; AAV38100 for SHV-1; AAP20891 for TEM-1; AEC32455 for TEM-52; BAM36530 for TEM-116; OBR18659 for KPC-2; AKN35351 for NMC-A; AAR06587 for SFC-1; CAA82281 for SME-1. The acylenzyme structures of these enzymes bound to meropenem were predicted using Protenix (Protenix-v1) [11, 12] by forming a covalent bond between Ser70 hydroxyl oxygen (TEM-1 numbering) and the electrophilic carbon of meropenem. Multiple sequence alignment and template features were enabled during the prediction. The sample number, recycle number, and diffusion steps were set to their default values. The predicted model with the highest overall pLDDT (predicted Local Distance Difference Test) score for each acylenzyme complex was selected for subsequent calculations.

#### *System setup and preparation*

The deacylating water (DW), which is crucial for antibiotic deacylation, was added manually to all predicted acylenzyme complexes by placing it between Glu166 and the electrophilic carbonyl carbon of meropenem, consistent with the location required for nucleophilic attack [13]. Force field parameters of Ser70-acylated meropenem were obtained from previous work [14]. In short, partial charges were calculated using RESP charge calculation based on HF/6-31G(d) of the capped Ser70-meropenem fragment using the R.E.D server [15], while atom types and bond parameters were described by GAFF force field [16]. The proteins were parameterized using the Amber ff14SB force field [17]. The protonation states of ionizable residues at pH 7.0 were determined using PropKa3.1 [18], and histidine tautomers were assigned for each acylenzyme complex individually using the reduce program from AmberTools24 [19]. All ionizable residues were predicted to adopt their standard protonation states, with Asp and Glu deprotonated, Lys and Arg protonated, and His singly protonated at either NE2 or ND1 atom. The only exception was Asp214 (TEM-1 numbering), which was predicted to be protonated in all enzymes. All acylenzyme complexes were solvated in a rectangular box of TIP3P water [20], with a minimum distance between the complex and the box edge of 10 Å. Each acylenzyme complex was neutralised by replacing random bulk water molecules with either $Na^+$ or $Cl^-$ ions as needed.

##### *Acylenzyme complex and approximate transition state (ATS) sampling*

All systems were minimized using 2500 steps of steepest descent followed by 2500 steps of conjugate gradient. The minimized acylenzyme structures were further optimized using quantum mechanics/molecular mechanics (QM/MM) minimization to generate reasonable acylzyme and ATS structures. The same steepest descent and conjugate gradient steps were used. The same QM region as that used in previous work [1] was applied. The QM region consists of the core structure of meropenem, the sidechains of Ser70 and the general base Glu166 (TEM-1 numbering), and the DW, resulting in a total of 44 atoms (including 3 link atoms) with a total charge of −2e (**Figure S2**). The QM region was described by the semi-empirical method DFTB2 (SCC-DFTB) [21] as used in previous work [1, 22]. To maintain reasonable and reactive conformations for both the acylenzyme and ATS, two one-sided weak distance harmonic restraints (10 kcal $mol^{-1}$ $Å^{-2}$ force constant) were applied between the meropenem carbonyl oxygen and Ser70 backbone amine, and between meropenem carbonyl carbon and oxygen, allowing free sampling when the distance is below 2.5 Å and 1.2 Å respectively. Three one-sided distance harmonic restraints (50 kcal $mol^{-1}$ $Å^{-2}$ force constant) were also applied on the DW O-H, meropenem hydroxyethyl O-H and Ser70 backbone N-H to maintain the distances below 1.2 Å. In addition, two reaction coordinates were used to generate ATS from MM minimized acylenzyme structures. One is proton transfer coordinate between Glu166 and DW: distance ($d$[Glu@OE2, DW@H] − $d$[DW@O, DW@H]) which shows the break of O-H bond in DW and formation of Glu@OE2-DW@H bond. The other is nucleophilic attack involves with the $OH^-$ of DW acting nucleophilic attack ($d$[DW@O, MER@C1]) to electrophilic carbon and forming a covalent bond. The approximate values for these two reaction coordinates were set to −0.1 Å and 1.7 Å respectively, based on previous free energy surfaces of meropenem deacylation by the same enzymes [1]. The force constant for both reaction coordinates was set to 5000 kcal $mol^{-1}$ $Å^{-2}$.

Then, all systems were heated from 50 K to 300 K over a period of 5 ps, with 3 replicas generated per system. All systems were further simulated in the NPT ensemble at 300 K and 1 bar for 25 ps or 7 ps, with first 5 ps discarded for analysis (as relaxation). Temperature was maintained using Langevin dynamics with a collision frequency of 0.2 $ps^{-1}$, while pressure was controlled using a Berendsen barostat with isotropic position scaling and a pressure relaxation time of 1 ps. The same QM region and harmonic restraints were used, except that the force constants for the reaction coordinates were set to 100 kcal $mol^{-1}$ $Å^{-2}$. Periodic boundary conditions were applied during all simulations and the SHAKE algorithm was applied to fix all bond lengths involving MM hydrogen atoms. No SHAKE algorithm was used in the QM region. The time step of the simulations was set to 0.1 fs, and the non-bonded interactions cutoff was set to 8.0 Å. The simulation trajectories were saved every 0.1 ps, resulting in 200 frames per replica (20 ps sampling) and a total of 600 frames per system for subsequent analysis and electric field calculations, or 20 frames per replica (2 ps sampling) and a total of 60 frames per system. All simulations were conducted using Amber24/AmberTools24 [19] and trajectories were analysed using CPPTRAJ [23].

##### *Electric field calculations*

The effective electric field ($E_{eff}$) was calculated as the projection of the electric field onto the C=O bond of the meropenem carbonyl group using the FieldTools [1]. Similarly, the same "reactive" part of the substrate as in the previous work [1], together with the entire hydroxyethyl group, was excluded from the calculations. The $E_{eff}$ standard deviation for each system was calculated from the three replica means, which were obtained by first averaging the corresponding values within each replica.

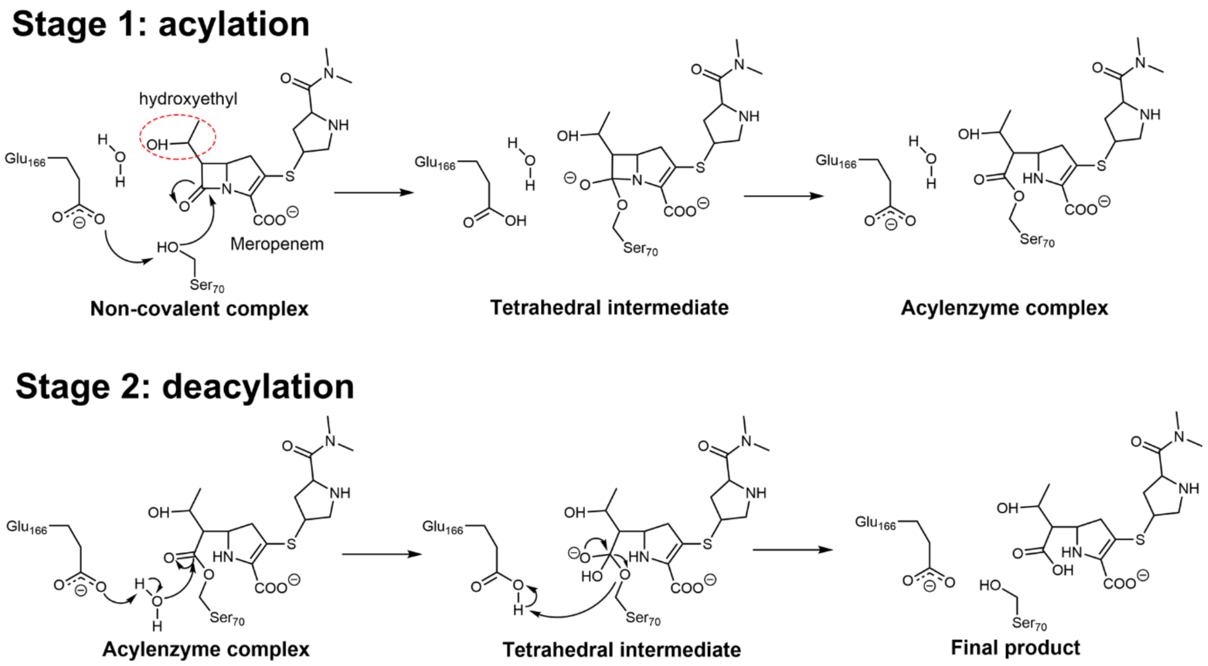


**Figure S1.** Reaction mechanism of meropenem hydrolysis by class A *β*-lactamases [24, 25]. The first half of deacylation is the conversion of the acylenzyme complex (AC) into the tetrahedral intermediate (TI), through Glu166-mediated activation of the deacylating water and subsequent nucleophilic attack on the acylenzyme carbonyl.

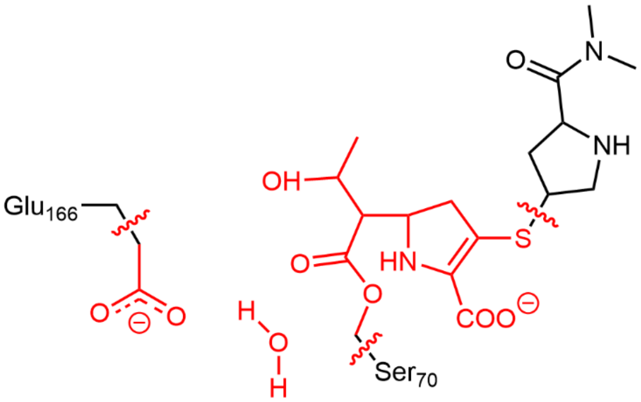

**Figure S2.** QM region in QM/MM (DFTB2/ff14SB) minimization and sampling. The atoms of QM region are coloured in red, and the breaks between QM and MM region are shown with wavy lines.

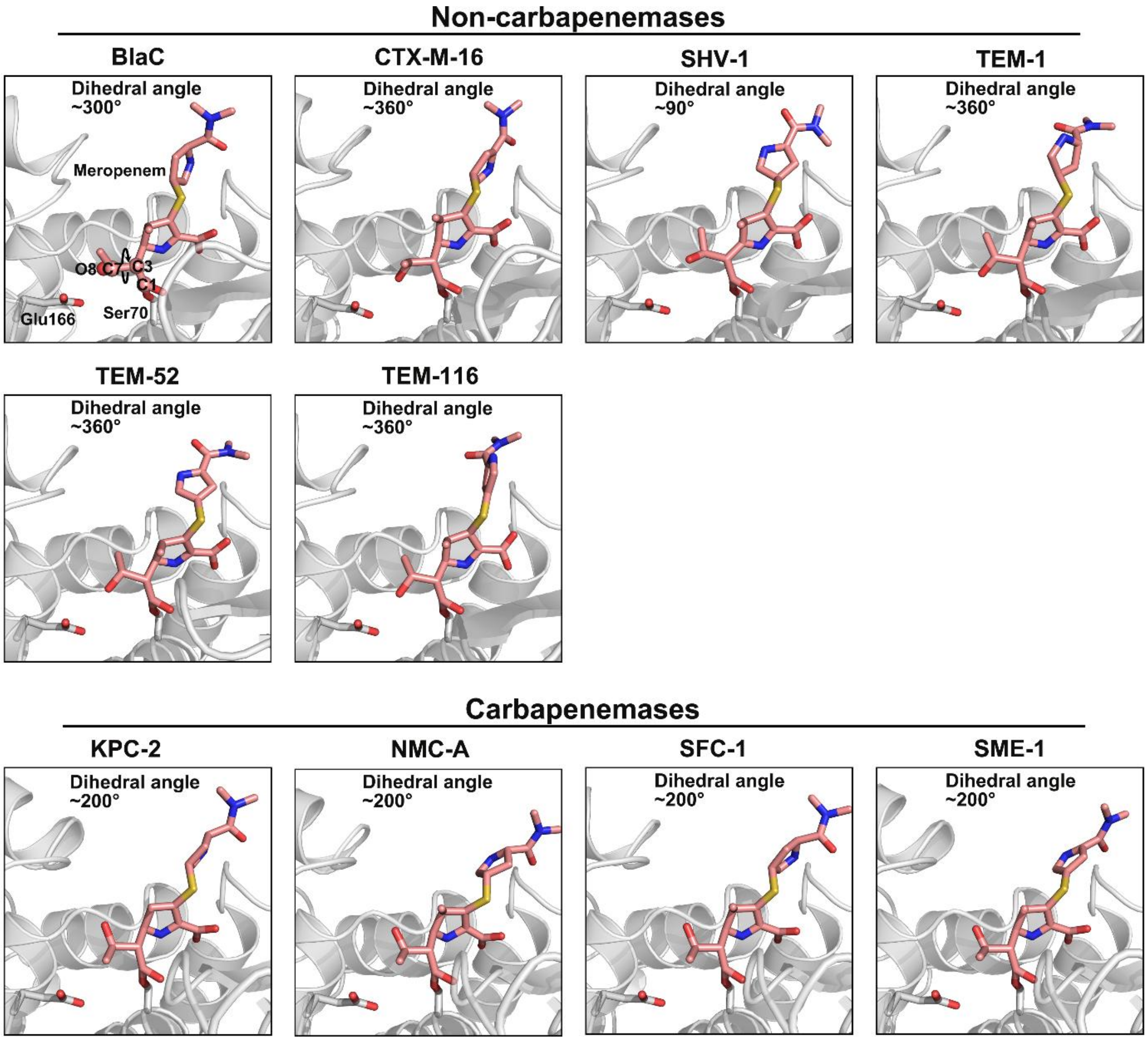


**Figure S3.** The Protenix predicted acylenzyme complexes of class A *β*-lactamases with meropenem. The catalytic residue Ser70 (gray), general base Glu166 (gray) and meropenem (pink) are shown as sticks. The atoms used to measure the hydroxyethyl group dihedral angle (C1-C3-C7-O8) are shown as spheres and labelled (first panel). The dihedral angle of hydroxyethyl group for each acylenzyme complex is labelled.

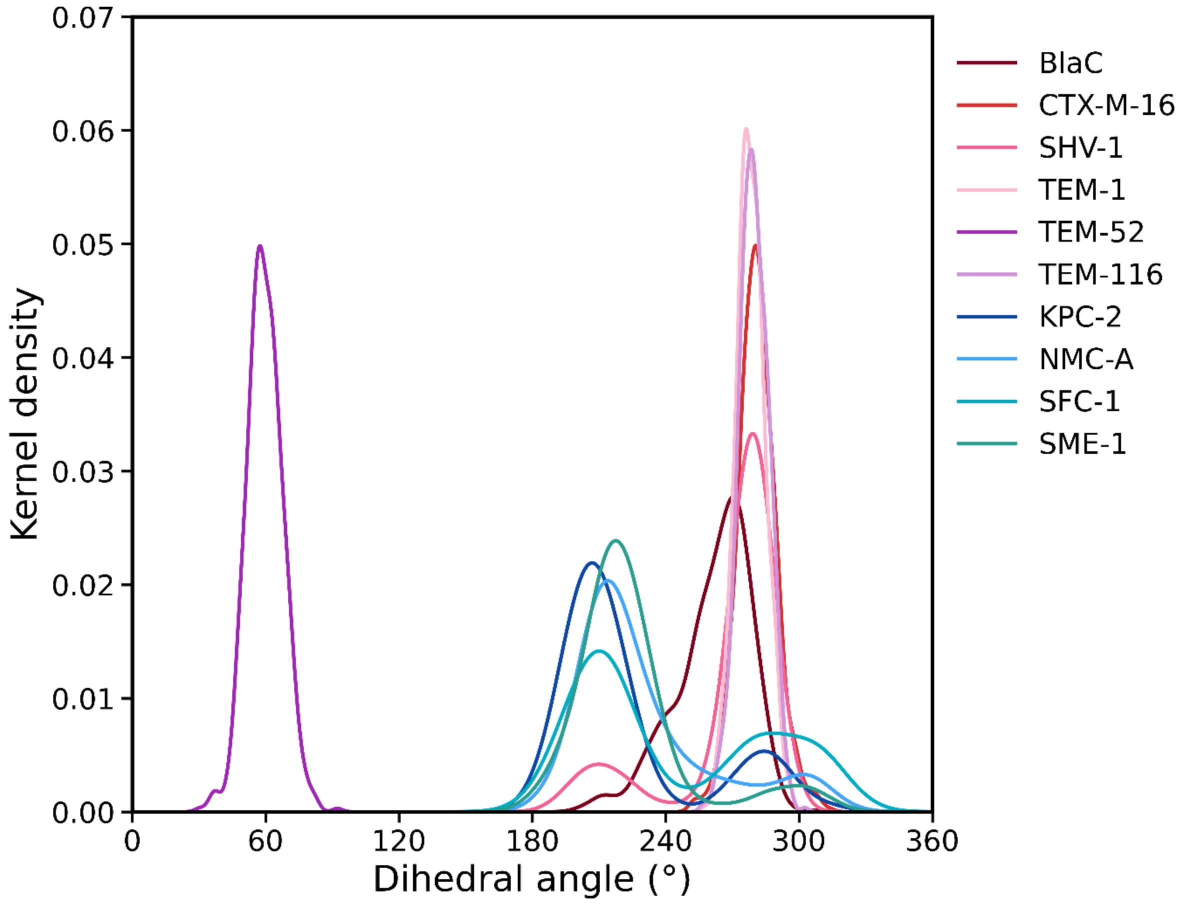


**Figure S4.** Distribution (kernel density estimate) of the meropenem hydroxyethyl group dihedral angle obtained from the QM/MM approximate transition state (ATS) sampling. Each distribution combines data from three independent replicas.

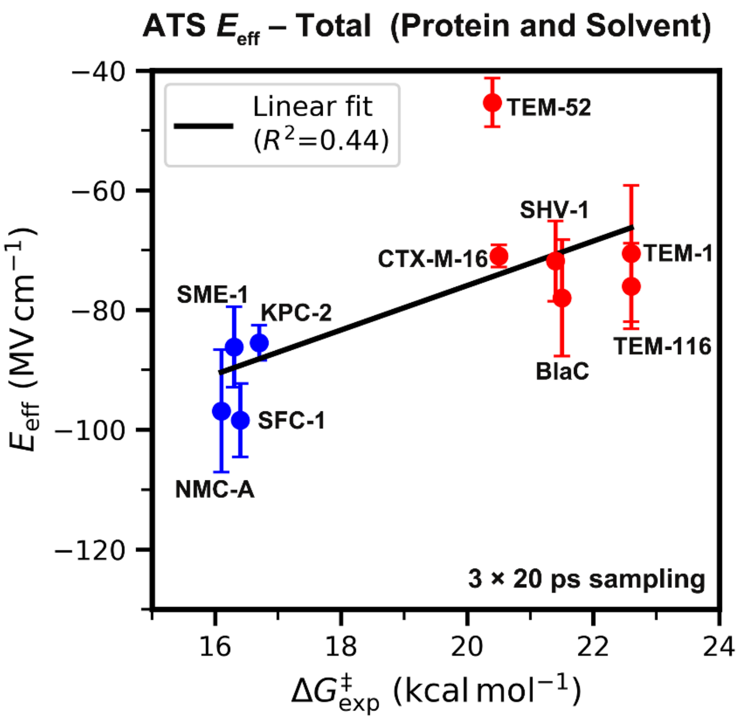


**Figure S5.** Calculated effective electric field (total $E_{eff}$, protein and solvent) along the meropenem C=O bond based on approximate transition state (ATS) sampling, with the hydroxyethyl group of meropenem included in the calculation. The correlation between $E_{eff}$ and the experimental energy barrier is shown. Carbapenemases are coloured in blue, whereas the non-carbapenemases are coloured in red. Error bars for $E_{eff}$ represent the standard deviation of the mean value across three replicas.

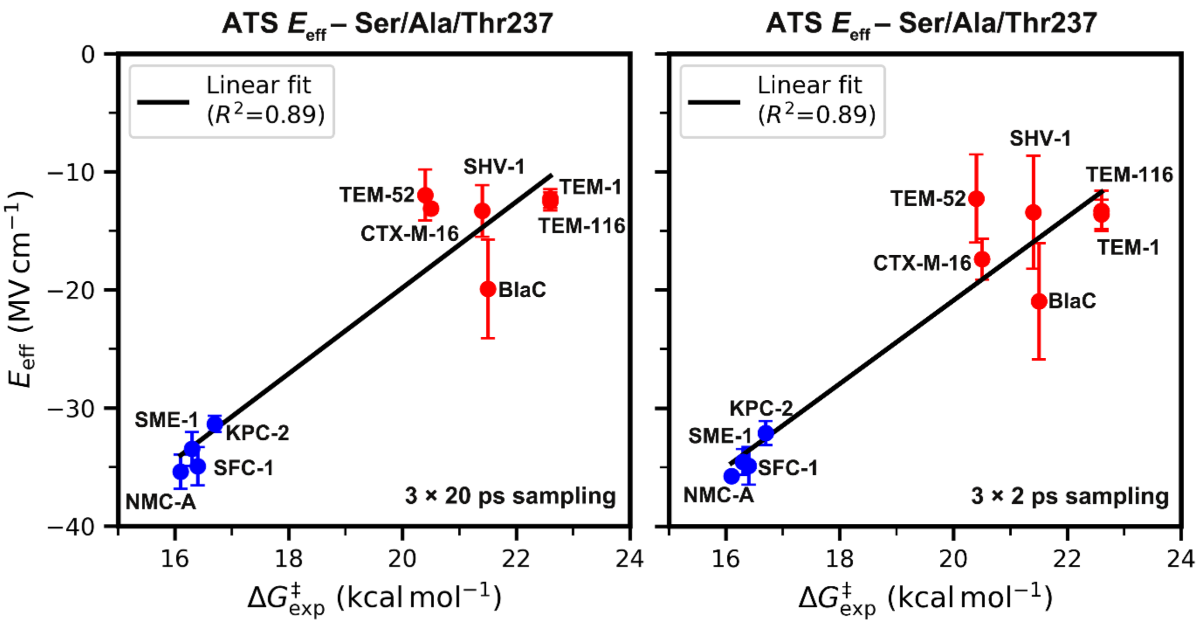


**Figure S6.** Calculated effective electric field ($E_{eff}$) contribution from the oxyanion-hole residue 237 (TEM-1 numbering) based on ATS sampling of 20 ps (left panel) and 2 ps (right panel) per replica. The correlation between $E_{eff}$ and experimental energy barrier is shown. Carbapenemases are coloured in blue, whereas the non-carbapenemases are coloured in red. Error bars for $E_{eff}$ represent the standard deviation of the mean value across three replicas.

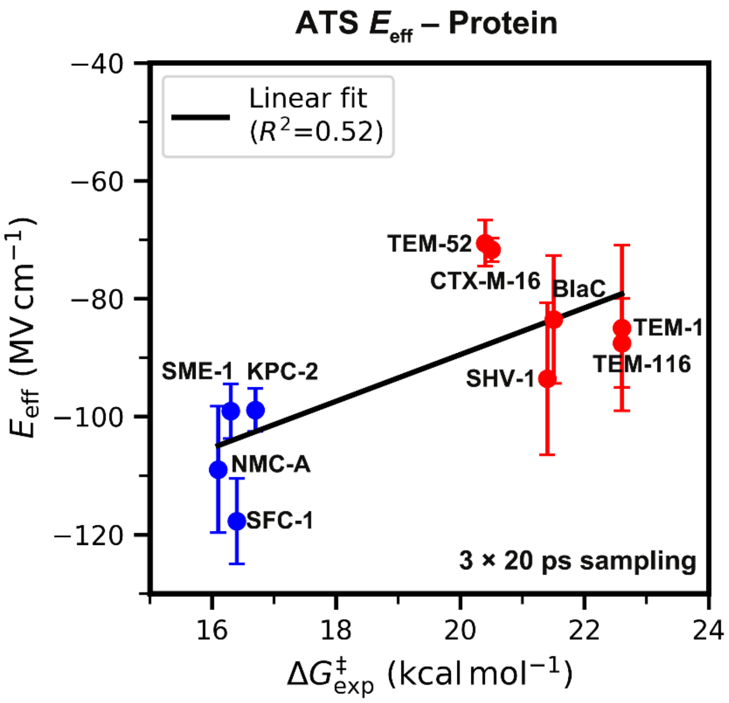


**Figure S7.** Calculated protein effective electric field ($E_{eff}$) along the meropenem C=O bond based on approximate transition state (ATS) sampling, with the hydroxyethyl group of meropenem excluded in the calculation. The correlation between $E_{eff}$ and experimental energy barrier is shown. Carbapenemases are coloured in blue, whereas the non-carbapenemases are coloured in red. Error bars for $E_{eff}$ represent the standard deviation of the mean value across three replicas.

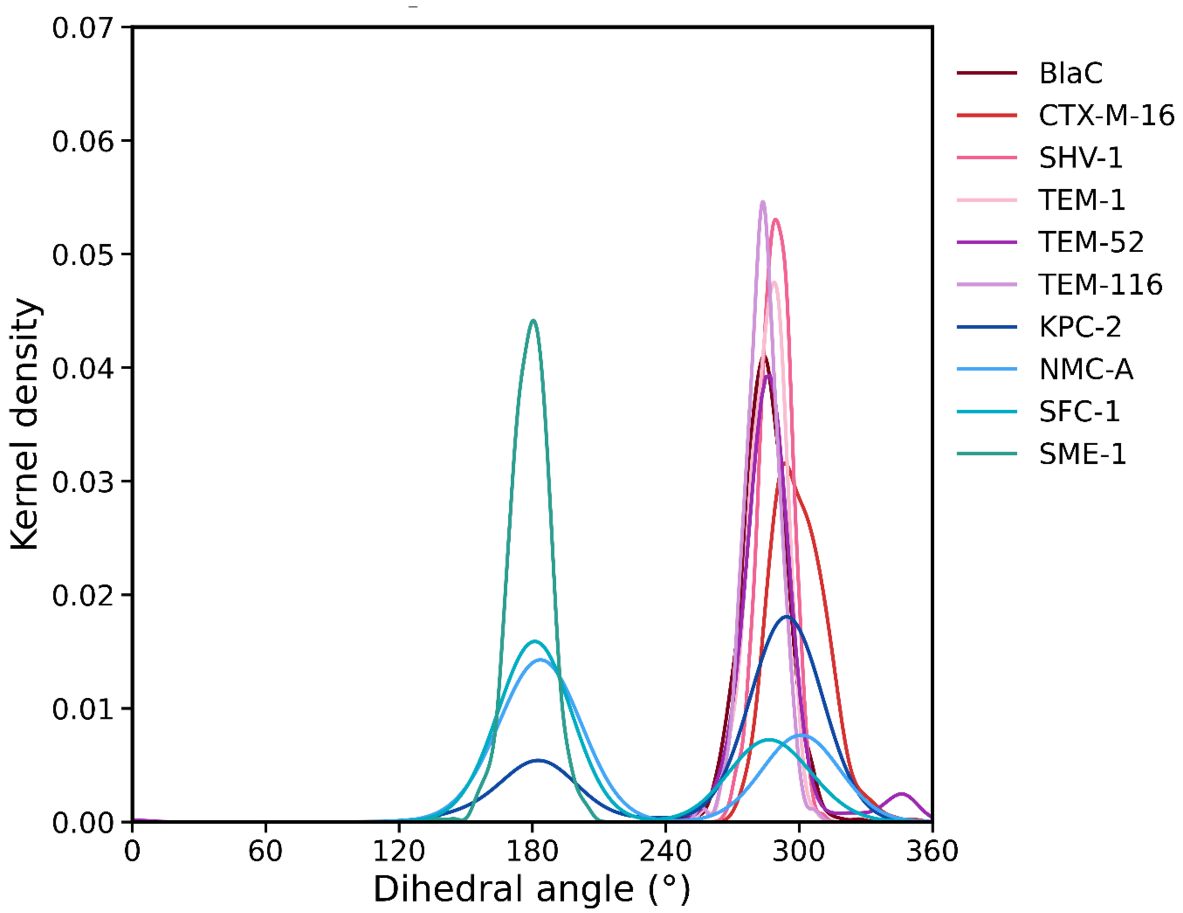


**Figure S8.** Distribution (kernel density estimate) of the meropenem hydroxyethyl group dihedral angle obtained from the QM/MM ayclenzyme complex sampling.

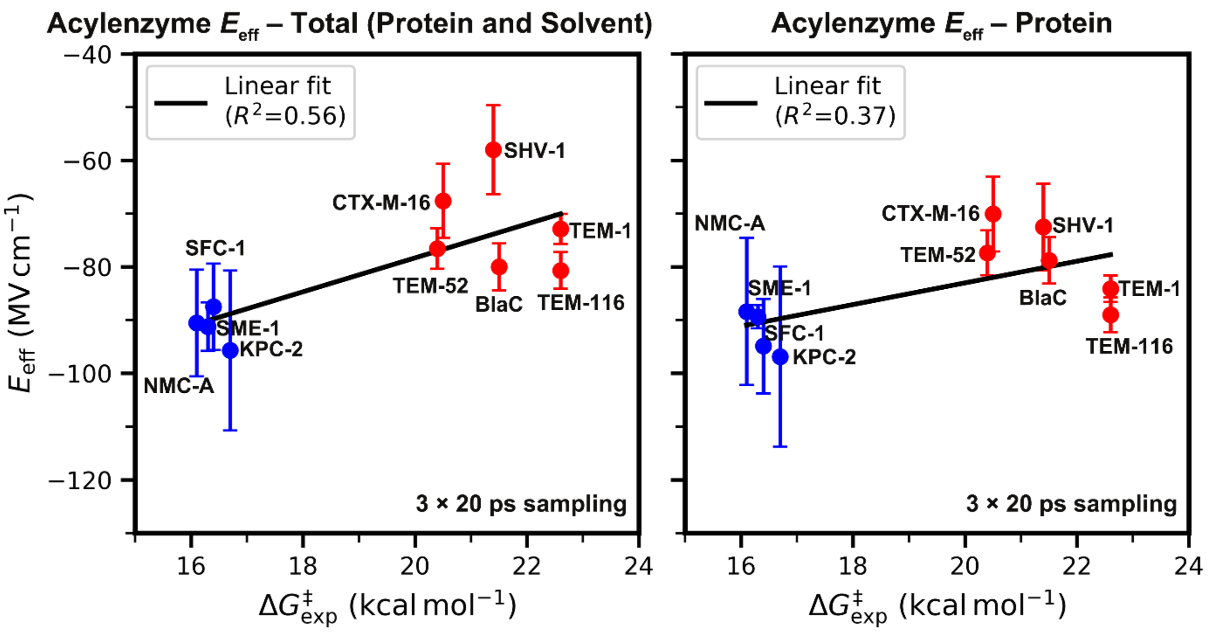


**Figure S9.** Calculated total (left panel) and protein (right panel) effective electric field ($E_{eff}$) along the meropenem C=O bond based on acylenzyme complex sampling, with the hydroxyethyl group of meropenem excluded in the calculation. The correlation between $E_{eff}$ and experimental energy barrier is shown. Carbapenemases are coloured in blue, whereas the non-carbapenemases are coloured in red. Error bars for $E_{eff}$ represent the standard deviation of the mean value across three replicas.

**Table S1. Calculated total and protein effective electric fields ($E_{eff}$, hydroxyethyl group excluded) for acylenzyme complexes and approximate transition state (ATS) ensembles (3 × 20 ps sampling per system)**

| Enzyme | Acylenzyme | | ATS | | $\Delta G^{\ddagger}_{exp}$ (kcal $mol^{-1}$)[b] |
|---|---|---|---|---|---|
| | Total $E_{eff}$ (MV $cm^{-1}$)[a] | Protein $E_{eff}$ (MV $cm^{-1}$)[a] | Total $E_{eff}$ (MV $cm^{-1}$)[a] | Protein $E_{eff}$ (MV $cm^{-1}$)[a] | |
| BlaC | −80.0 ± 4.4 | −78.8 ± 4.4 | −83.2 ± 8.5 | −83.5 ± 10.9 | 21.5[c] |
| CTX-M-16 | −67.6 ± 7.0 | −70.1 ± 7.0 | −71.7 ± 1.3 | −71.7 ± 2.0 | 20.5[d] |
| SHV-1 | −58.0 ± 8.4 | −72.5 ± 8.1 | −76.2 ± 11.2 | −93.6 ± 12.9 | 21.4[e] |
| TEM-1 | −72.9 ± 2.8 | −84.1 ± 2.5 | −71.9 ± 11.7 | −85.0 ± 14.0 | 22.6[e] |
| TEM-52 | −76.6 ± 3.8 | −77.4 ± 4.2 | −66.6 ± 4.3 | −70.6 ± 3.9 | 20.4[f] |
| TEM-116 | −80.7 ± 3.4 | −89.0 ± 3.3 | −76.8 ± 7.2 | −87.5 ± 7.5 | 22.6[e] |
| KPC-2 | −95.7 ± 15.0 | −96.9 ± 16.9 | −98.3 ± 2.9 | −98.9 ± 3.6 | 16.7[g] |
| NMC-A | −90.5 ± 10.0 | −88.4 ± 13.8 | −108.1 ± 10.9 | −109.0 ± 10.7 | 16.1[h] |
| SFC-1 | −87.5 ± 8.1 | −94.9 ± 8.9 | −108.5 ± 4.8 | −117.7 ± 7.2 | 16.4[i] |
| SME-1 | −91.2 ± 4.5 | −89.4 ± 2.2 | −97.9 ± 7.0 | −99.1 ± 4.6 | 16.3[j] |

[a] Standard deviations obtained from three independent simulations.

[b] Experimental free energy barriers are taken from a previous study [1].

[c] $k_3$ value is from Chow *et al* [8].

[d] The $k_{cat}$ value of CTX-M-14, which is a close variant to CTX-M-16 with comparable activity spectrum is used here. The $k_{cat}$ value is from Ishii *et al* [10].

[e] $k_{cat}$ value of TEM-1 is from Queenan *et al* [6]. TME-116 only have two point mutations compared with TEM-1, and these mutations are related to the stability of enzyme instead of activity. Therefore, the same $k_{cat}$ value is used for TEM-116.

[f] $k_3$ value is from Franceschini *et al* [9]. Noting that acylation rather than deacylation is rate-limiting for TEM-52 [9].

[g] $k_{cat}$ value is calculated as the average of the $k_{cat}$ values reported in three references [2, 6, 7].

[h] $k_{cat}$ value is from MariotteBoyer *et al* [4].

[i] $k_{cat}$ value is from Fonseca *et al* [3].

[j] $k_{cat}$ value is from Queenan *et al* [5].

**Table S2. Availability of carbapenem acylenzyme complexes with studied class A *β*-lactamases in the PDB database before the Protenix-v1 training set cutoff (30 September 2021)**

| Acylenzyme | | PDB ID |
|---|---|---|
| Carbapenem | Enzyme | |
| Meropenem | BlaC | 3DWZ [26] |
| | SHV-1 | 2ZD8 [27] |
| | SFC-1 E166A | 4EV4 [28] |
| | SFC-1 S70A | 4EUZ [28] |
| Imipenem | TEM-1 N132A | 1JVJ [29] |
| | TEM-116 | 1BT5 [30] |
| | KPC-2 F72Y | 7LLH [31] |
| | SME-1 | 1DY6 [32] |

(30) Maveyraud, L.; Mourey, L.; Kotra, L. P.; Pedelacq, J. D.; Guillet, V.; Mobashery, S.; Samama, J. P. Structural basis for clinical longevity of carbapenem antibiotics in the face

of challenge by the common class A $\beta$-lactamases from the antibiotic-resistant bacteria. *J Am Chem Soc* **1998**, *120* (38), 9748-9752. DOI: 10.1021/ja9818001.

(31) Furey, I. M.; Mehta, S. C.; Sankaran, B.; Hu, L.; Prasad, B. V. V.; Palzkill, T. Local interactions with the Glu166 base and the conformation of an active site loop play key roles in carbapenem hydrolysis by the KPC-2 $\beta$-lactamase. *J Biol Chem* **2021**, *296*, 100799. DOI: 10.1016/j.jbc.2021.100799.

(32) Sougakoff, W.; L'Hermite, G.; Pernot, L.; Naas, T.; Guillet, V.; Nordmann, P.; Jarlier, V.; Delettré, J. Structure of the imipenem-hydrolyzing class A $\beta$-lactamase SME-1 from *Serratia marcescens*. *Acta Crystallogr D* **2002**, *58*, 267-274. DOI: 10.1107/S0907444901019606.
